# *Yersinia pseudotuberculosis* YopE prevents uptake by M cells and instigates M cell extrusion in human ileal enteroid-derived monolayers

**DOI:** 10.1101/2021.08.23.457340

**Authors:** Alyssa C. Fasciano, Gaya S. Dasanayake, Mary K. Estes, Nicholas C. Zachos, David T. Breault, Ralph R. Isberg, Shumin Tan, Joan Mecsas

## Abstract

Many pathogens use M cells to access the underlying Peyer’s patches and spread to systemic sites via the lymph as demonstrated by ligated loop murine intestinal models. However, the study of interactions between M cells and microbial pathogens has stalled due to the lack of cell culture systems. To overcome this obstacle, we use human ileal enteroid-derived monolayers containing five intestinal cell types including M cells to study the interactions between the enteric pathogen, *Yersinia pseudotuberculosis* (*Yptb*) and M cells. The *Yptb* type three secretion system (T3SS) effector Yops inhibit host defenses including phagocytosis and are critical for colonization of the intestine and Peyer’s patches. Therefore, it is not understood how *Yptb* traverses through M cells to breach the epithelium. By growing *Yptb* under two physiological conditions that mimic the early infectious stage (low T3SS-expression) or host-adapted stage (high T3SS-expression), we found that large numbers of *Yptb* specifically associated with M cells, recapitulating murine studies. Transcytosis through M cells was significantly higher by *Yptb* expressing low levels of T3SS, because YopE and YopH prevented *Yptb* uptake. YopE also caused M cells to extrude from the epithelium without inducing cell-death or disrupting monolayer integrity. Sequential infection with early infectious stage *Yptb* reduced host-adapted *Yptb* association with M cells. These data underscore the strength of enteroids as a model by discovering that Yops impede M cell function, indicating that early infectious stage *Yptb* more effectively penetrates M cells while the host may defend against M cell penetration of host-adapted *Yptb*.

## Introduction

Many diverse bacterial and viral pathogens, including enteric *Yersinia* spp., exploit specialized epithelial microfold (M) cells that reside within the follicular associated epithelium (FAE) above Peyer’s patches (PP) to gain access to deeper tissues.^1–3^ M cells play a major role in immunosurveillance by taking up and delivering luminal contents to PP and have distinct morphology including apical polarization of β1 integrin and the lack of tightly packed apical microvilli present on neighboring enterocytes.^1,4–6^ The enteropathogenic bacteria, *Yersinia pseudotuberculosis* (*Yptb)* and *Y. enterocolitica*, are primarily transmitted via ingestion of contaminated food by the fecal-oral route and are psychrotrophs that grow at temperatures below 4°C.^7^ Thus, outbreaks have been associated with food stored in the cold such as milk products and vegetables.^8–12^ After ingestion, *Yptb* primarily target to the terminal ileum and gain access to underlying lymph tissues to cause acute intestinal illness and mesenteric lymphadenitis and in rare cases can spread systemically.^13^ In murine intestinal ligated loop infection models, *Yptb* has been shown to bind to β1 integrins on M cells using the bacterial adhesin protein invasin.^2,14^ After breaching the intestinal layer, *Yersinia* establish infection in PP and form extracellular microcolonies derived from a clonal bacterium.^15–18^

Critical to *Yptb* virulence is a type three secretion system (T3SS) encoded on a 70kb plasmid that functions to inject effector proteins, called Yops, into host cells disrupting antimicrobial functions.^19–22^ T3SS expression is regulated by temperature and is induced at 37°C ^23,24^, so *Yptb* consumed via food in cold-storage conditions may reach the intestine before expressing the T3SS. When *Yptb* express the T3SS, host cell contact, mediated by adhesin proteins invasin and YadA, induces Yop injection directly into the host cell cytoplasm.^25,26^ YopE, YopH, and YopO contribute to the anti-phagocytic activity of *Yptb.*^27–31^ YopE is a GTPase-activating protein (GAP) for the Rho family of GTP-binding proteins.^28,32^ It prevents *Yptb* internalization by non-polarized epithelial cells via its Rac1-GAP activity and causes cell rounding in non-polarized epithelial cells and limits Yop injection via its RhoA-GAP activity.^28,33,34^ YopH is a protein tyrosine phosphatase and contributes to inhibiting phagocytosis by immune cells and uptake by epithelial cells.^27,35,36^ YopO has serine/threonine kinase and RhoGDI activities and *Y. enterocolitica* YopO participates in preventing internalization by macrophages and neutrophils *in vitro.*^37,38^

YopE and YopH are required for *Yptb* survival in intestinal tissues and PP ^39,40^, yet paradoxically function to prevent internalization raising the conundrum of how *Yptb* traverses across the FAE containing M cells to establish infection. The attenuation of *yopE* and *yopH* mutants in the GI tract ^39,40^ and the sparsity of M cells in the epithelium has made it difficult to study the detailed molecular host-pathogen interactions that occur at the host epithelium using murine models. So whether *Yptb* Yops are delivered into and/or function within M cells has not been described. Further, widely used *in vitro* cell culture systems do not mimic the microenvironment and cellular complexity of the human intestine.^41^ Using human enteroids derived from intestinal biopsies *in vitro* has helped progress the study of intestinal infections (reviewed in ^42,43^), with a key advantage being the ability of self-renewing Lgr5+ intestinal stem cells to differentiate into four major intestinal cell subpopulations. ^44,45^ While these cultures initially lack M cells, we and others have described a method to induce development of functional M cells in human ileal enteroids by adding RANKL and TNFα to differentiation media. ^46–49^

In the current study, we used human ileal enteroid-derived monolayers containing M cells to reconstruct the cellular microenvironment of the intestine and to visualize how *Yptb* exploit M cells to gain access to underlying lymph tissues. We find that the consequences of the interaction between M cells and *Yptb* depend on T3SS status. *Yptb* lacking T3SS expression more efficiently breached monolayers containing M cells and reduced the association of T3SS-expressing *Yptb* with M cells in subsequent infection. Conversely, T3SS-expressing *Yptb* prevented M cell invasion and induced M cell extrusion indicating methods by which infection in PP may be limited. Thus, these studies with enteroids contribute new visual and mechanistic insights into early interactions between *Yptb* and M cells in the human intestinal epithelium that were previously unattainable in other models.

## Results

### Large numbers of *Yptb* bind to enteroid-derived M cells in an invasin-dependent manner

Infection of murine ligated loop models have established that *Yptb* target to M cells. However, it has been challenging to understand cellular events at the *Yptb*-M cell interface due to the lack of an appropriate intestinal cell culture system that faithfully recapitulates the cellular microenvironment found in the ileum overlaying PP. We overcome this critical block in the field by using human ileal enteroid-derived monolayers containing at least four major intestinal cell types and M cells.^49^ To investigate the ability of *Yptb* to bind to these intestinal cells, monolayers derived from human ileal enteroid line HIE25 were differentiated in the presence of RANKL and TNFα (RT+), conditions that promote M cell development ^49^, or in their absence (RT-). Monolayers were apically infected with WT *Yptb* YPIII expressing green fluorescent protein (GFP). After 5-hour infection, RT- monolayers had few *Yptb* bound to the apical surface of cells (Fig 1A, white circles). Investigation of the orthogonal planes indicated that a single *Yptb* bacterium was often located at intercellular junctions (Fig 1B, orange arrows). By contrast, RT+ monolayers had many *Yptb* bound to the surface of GP2^+^ M cells (Fig 1C-D, green arrows). Quantification of *Yptb* binding showed that more than 90% of *Yptb*-associated cells in RT- monolayers had only 1 *Yptb* bound per cell, while the remaining cells had 2-5 *Yptb* bound (Fig 1E). In RT+ monolayers, over 40% of *Yptb*-associated cells had 6+ *Yptb* bound and these cells often had over 30 *Yptb* bound per cell (Fig 1C-E). The remaining cells had either 1 *Yptb* or 2-5 *Yptb* bound, consistent with the pattern of binding observed in RT- monolayers (Fig 1E).

**Figure 1.**
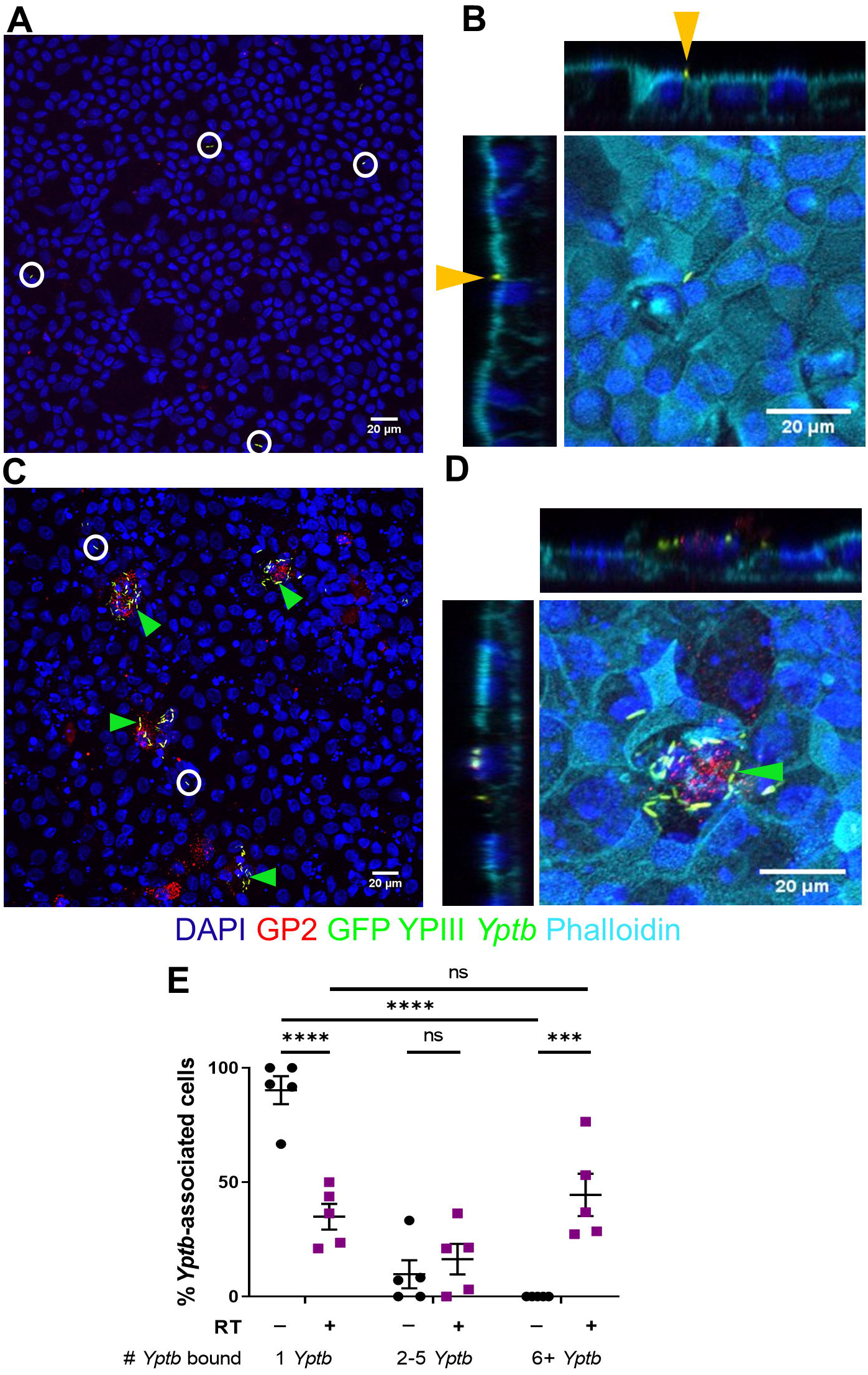
WT *Yptb* binds to enteroid-derived M cells in large numbers. HIE25 ileal monolayers were differentiated under (A-B) RT- or (C-D) RT+ conditions (200 ng/ml RANKL and 50 ng/ml TNFα). Monolayers were infected for 5 hours with 5×10^6^ CFU WT YPIII *Yptb* expressing GFP (green) and stained with anti-GP2 antibody (M cells-red), DAPI (nuclei-blue), and phalloidin (F-actin-cyan). XY planes are maximum intensity projections. White circles/orange arrows denote a single bacterium. Green arrows denote multiple *Yptb.* (B, D) Orthogonal XZ and YZ planes are shown. (E) The number of *Yptb* bound to a cell was counted, binned into groups of 1, 2-5, or 6+ *Yptb,* and the percentage of cells with the specified number of *Yptb* bound was plotted. Each point represents one Transwell. Bars indicate mean and SEM. Data were pooled from 3+ independent experiments with 2-4 fields analyzed per Transwell and averaged. Statistical significance was determined using a two-way ANOVA with Tukey’s post hoc multiple comparison test.

In RT+ monolayers, we occasionally noted that cells with 6+ *Yptb* bound had no or low expression of GP2 and that some GP2^+^ cells did not have *Yptb* bound (Fig 1C). Since GP2 is a late marker of M cells ^50,51^, we evaluated whether *Yptb* may be binding to different subsets of M cells by locating M cells with apical β1 integrin expression (Fig 2). Apical β1 integrin staining was detected in RT+ monolayers, but not in RT- monolayers (Fig 2A, B). As observed previously with GP2^+^ M cells ^49^, staining with the F-actin labeling dye phalloidin showed reduced apical F-actin expression on β1 integrin^+^ M cells compared to neighboring non-M cells (Fig 2C, D). These M cells were frequently bound by more than 6 *Yptb* (Fig 2E). Since no instances of 6+ *Yptb* bound to a cell were found in RT- monolayers, M cells were defined as cells with 6+ *Yptb* bound for further analyses.

**Figure 2.**
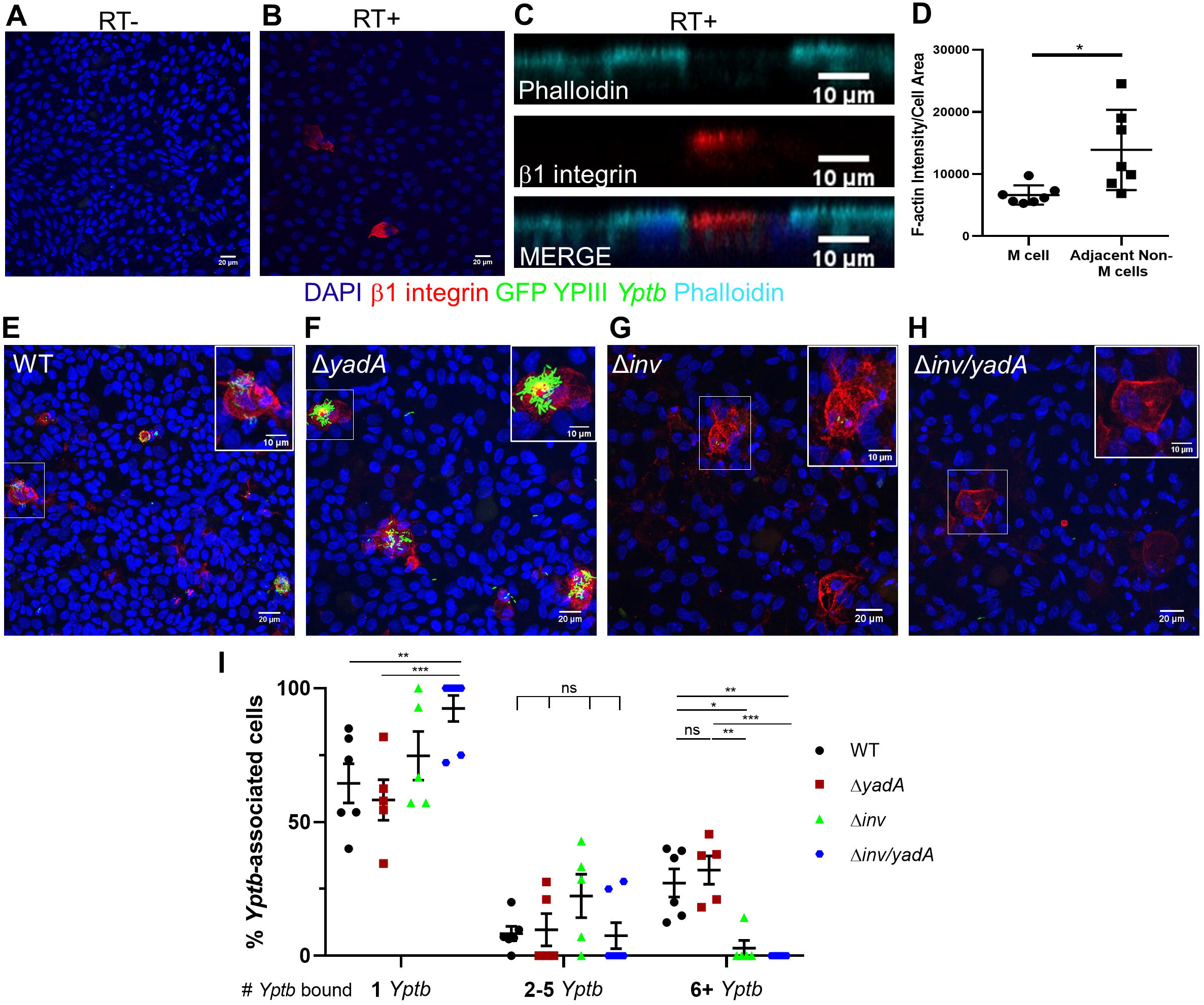
*Yptb* bind to M cells using invasin. (A-B) Differentiated uninfected HIE25 ileal monolayers (A) RT- or (B) RT+ were stained for apical β1 integrin (red) and DAPI (nuclei-blue). (C) XZ plane of RT+ monolayer shows β1 integrin^+^ M cell and neighboring non-M cells stained with phalloidin (F-actin-cyan). (D) Fluorescence intensity of F-actin was measured for each M cell and corresponding adjacent non-M cells and divided by cell area. Each point represents an M cell or the average of the corresponding neighboring non-M cells. Bars indicate mean and SD. Data were pooled from 3 images from 3 independent experiments. (E-I) Differentiated HIE25 RT+ monolayers were infected for 5 hours with 5×10^6^ CFU (E) WT, (F) Δ*yadA,* (G) Δ*inv,* or (H) Δ*inv/yadA* YPIII *Yptb* expressing GFP (green) and stained with anti-β1 integrin antibody (red) and DAPI (nuclei-blue). Magnified insets of an M cell are shown in upper right corner. XY planes are maximum intensity projections. (I) The number of *Yptb* bound to a cell was counted (1, 2-5, or 6+ *Yptb*) and the percentage of cells with the specified number of *Yptb* bound was plotted. Each point represents one Transwell. Bars indicate mean and SEM. Data were pooled from 3+ independent experiments with 2-4 fields analyzed per Transwell and averaged. Statistical significance was determined using a one-way ANOVA with Tukey’s post hoc multiple comparison test, with comparisons between columns within each row.

To determine which adhesin proteins are important for binding to M cells, RT+ monolayers were infected with WT *Yptb* and the adhesin mutant strains, Δ*yadA*, Δ*inv*, and Δ*inv/yadA,* all expressing GFP. There was no difference in the percentage of cells with 6+ *Yptb* bound when infected with WT and Δ*yadA* (Fig 2E, F, I). In the absence of invasin, almost no cells were detected with 6+ *Yptb* bound (Fig 2G, H, I). All strains bound singly or in smaller groups (1-5 bacteria) throughout the monolayers (Fig 2I). Thus, invasin promotes *Yptb* adhesion to enteroid-derived M cells.

Combined, these data show that *Yptb* bound sparsely to ileal epithelial monolayers, however when M cells were present, most bacteria were bound to M cells in large numbers in an invasin-dependent manner. Therefore, the enteroid-derived monolayer model is a valuable system that recapitulates *Yptb* binding observed *in vivo* in murine models. Importantly, this allows us for the first time to examine the fate of *Yptb*-infected M cells in the context of an epithelium that maintains distinct cellular functions.

### T3SS non-expressing *Yptb* efficiently gain access to the basolateral side in the presence of M cells compared to *Yptb* expressing the T3SS

To determine if *Yptb* reaches the basolateral side during infection in an M cell-dependent manner, HIE25 ileal monolayers were infected with WT *Yptb* grown at 37°C (WT 37°C), conditions which induce T3SS expression. Infections lasted 3 hours to limit bacterial replication in the basolateral chamber. Increased colony forming units (CFU) were detected in the basolateral chamber of RT+ monolayers compared to RT- monolayers (Fig 3A). The transepithelial electrical resistance (TEER) was taken to assess monolayer integrity and confirmed that the increase in CFU was not due to compromised barrier function in RT+ monolayers (TEER, 550 ± 57 Ω·cm^2^) compared to RT- monolayers (TEER, 619 ± 45 Ω·cm^2^).

**Figure 3.**
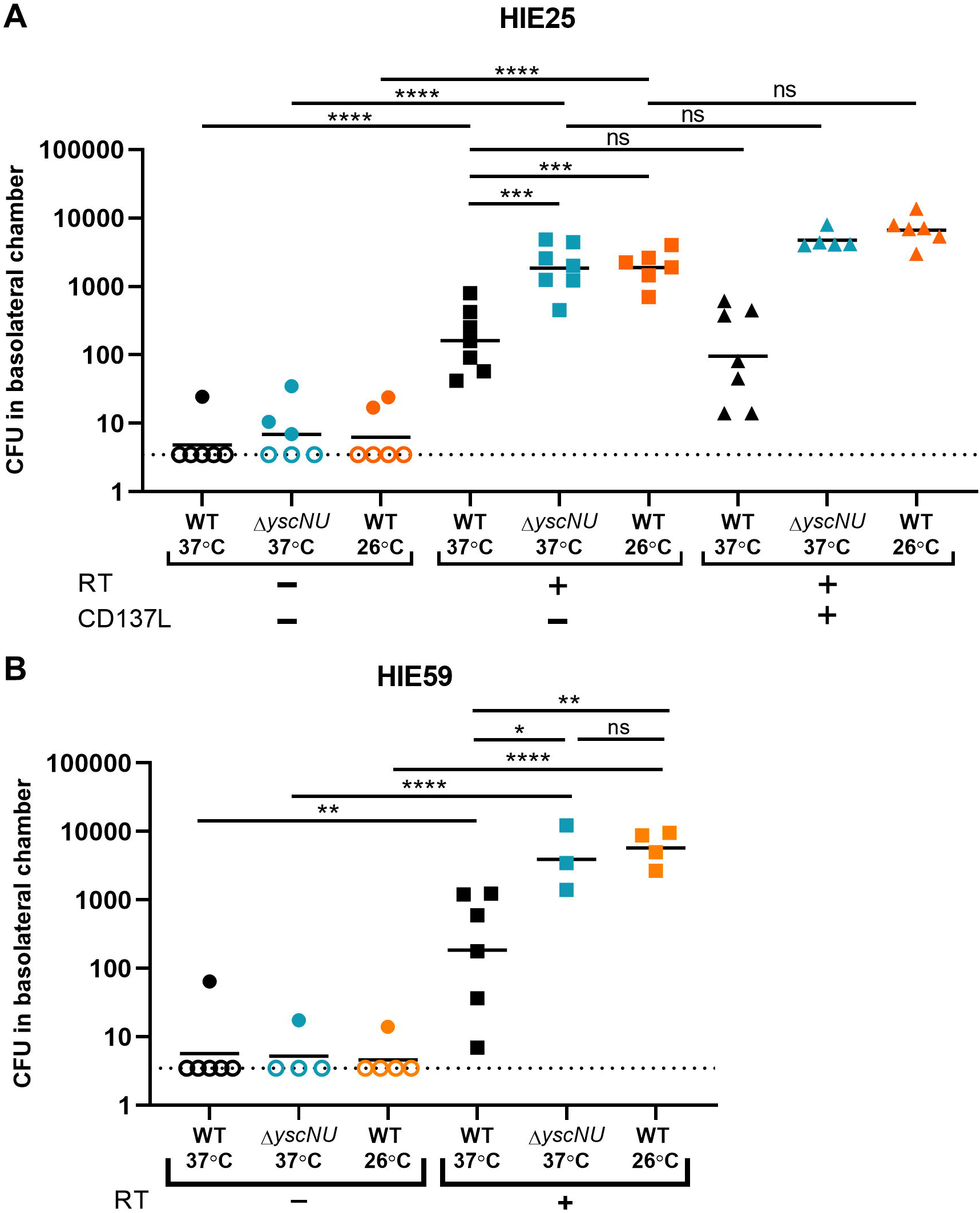
*Yptb* breaches the epithelium in the presence of M cells. (A) Differentiated HIE25 ileal monolayers were RT-, RT+, or RT+ with 100 ng/mL CD137L. Monolayers were infected for 3 hours with 5×10^6^ CFU of WT 37°C, Δ*yscNU* 37°C, or WT 26°C, all expressing GFP, and basolateral media was plated for CFU. Data were pooled from 3+ independent experiments. (B) Differentiated HIE59 ileal monolayers (RT- or RT+) were infected as in (A). Data were pooled from 2+ independent experiments. (A-B) Each point represents one Transwell. Bars indicate geometric mean. Dotted line indicates limit of detection. Open shapes indicate CFU values below limit of detection. Statistics were performed on the log-transformed values with a one-way ANOVA and Tukey’s post hoc multiple comparison tests.

Since the *Yptb* T3SS prevents internalization and phagocytosis ^19,30^, we evaluated the role of the T3SS in transcytosis by M cells using a strain of *Yptb* genetically lacking the T3SS (Δ*yscNU* 37°C) and WT *Yptb* grown at 26°C (WT 26°C), conditions under which the T3SS is not expressed.^24^ These strains were transported through M cell-containing monolayers at levels nearly 10-fold above WT 37°C, indicating that the absence of T3SS expression allows for more efficient access to the basolateral side. The CFU recovered for Δ*yscNU* 37°C and WT 26°C in RT- monolayers were near the limit of detection and similar to WT 37°C.

The receptor CD137 on M cells in mice has been reported to contribute to M cell functional maturity and increased transcytosis of particles.^52^ CD137L is the ligand for CD137 and is a TNF superfamily member that is present on stromal cells and bone marrow-derived cells in PP.^52^ To determine if CD137L increased enteroid-derived M cell functionality by means of increased transcytosis of *Yptb*, CD137L was added to the M cell induction media with RANKL and TNFα during differentiation. The CFU recovered in the basolateral chamber of RT+/CD137L monolayers were slightly but not significantly increased for the T3SS non-expressing strains with respect to RT+ monolayers (Fig 3A).

To confirm that M cell transcytosis was not confined to a single line of ileal enteroids, transcytosis was investigated with a different human ileal enteroid line, HIE59.^53^ As with HIE25, the CFU recovered in the basolateral chamber of Δ*yscNU* 37°C or WT 26°C infected RT+ monolayers were increased 10-fold compared to WT 37°C (Fig 3B). The CFU recovered in RT- monolayers were near or at the limit of detection in all infection conditions. Similarly, the TEER did not differ significantly between RT- (TEER, 681 ± 153 Ω·cm^2^) and RT+ (TEER, 588 ± 133 Ω·cm^2^) monolayers.

### *Yptb* T3SS reduces entry of *Yptb* into M cells

To gain insight into the result that T3SS non-expressing *Yptb* reached the basolateral compartment in larger numbers compared to *Yptb* expressing the T3SS, we analyzed the orthogonal planes of infected monolayers (Fig 4A-J). Most WT 37°C *Yptb* remained extracellularly bound to the surface of M cells, although uptake of a few *Yptb* by some M cells was observed (Fig 4B). To quantify the degree of bacterial internalization, we determined the percentage of M cells with 0 *Yptb* internalized (*Yptb* remained bound to the M cell surface), 1-9 *Yptb* internalized (*Yptb* on surface and internalized), or 10^+^ *Yptb* internalized (*Yptb* fully internalized) (Fig 4K, Table S1). About 80% of M cells infected with WT 37°C had no *Yptb* internalized and a smaller percentage of M cells had 1-9 *Yptb* internalized (Fig 4B, K, Table S1). By contrast, over 95% of M cells infected with Δ*yscNU* or WT 26°C had at least one *Yptb* internalized with over 80% of M cells having 10^+^ *Yptb* internalized (Fig 4C, D, K, Table S1). Combined with the transcytosis data, these results indicate that T3SS non-expressing *Yptb* are more efficiently internalized whereas T3SS expression greatly reduces, but does not fully prevent, *Yptb* internalization and transcytosis by M cells.

**Figure 4.**
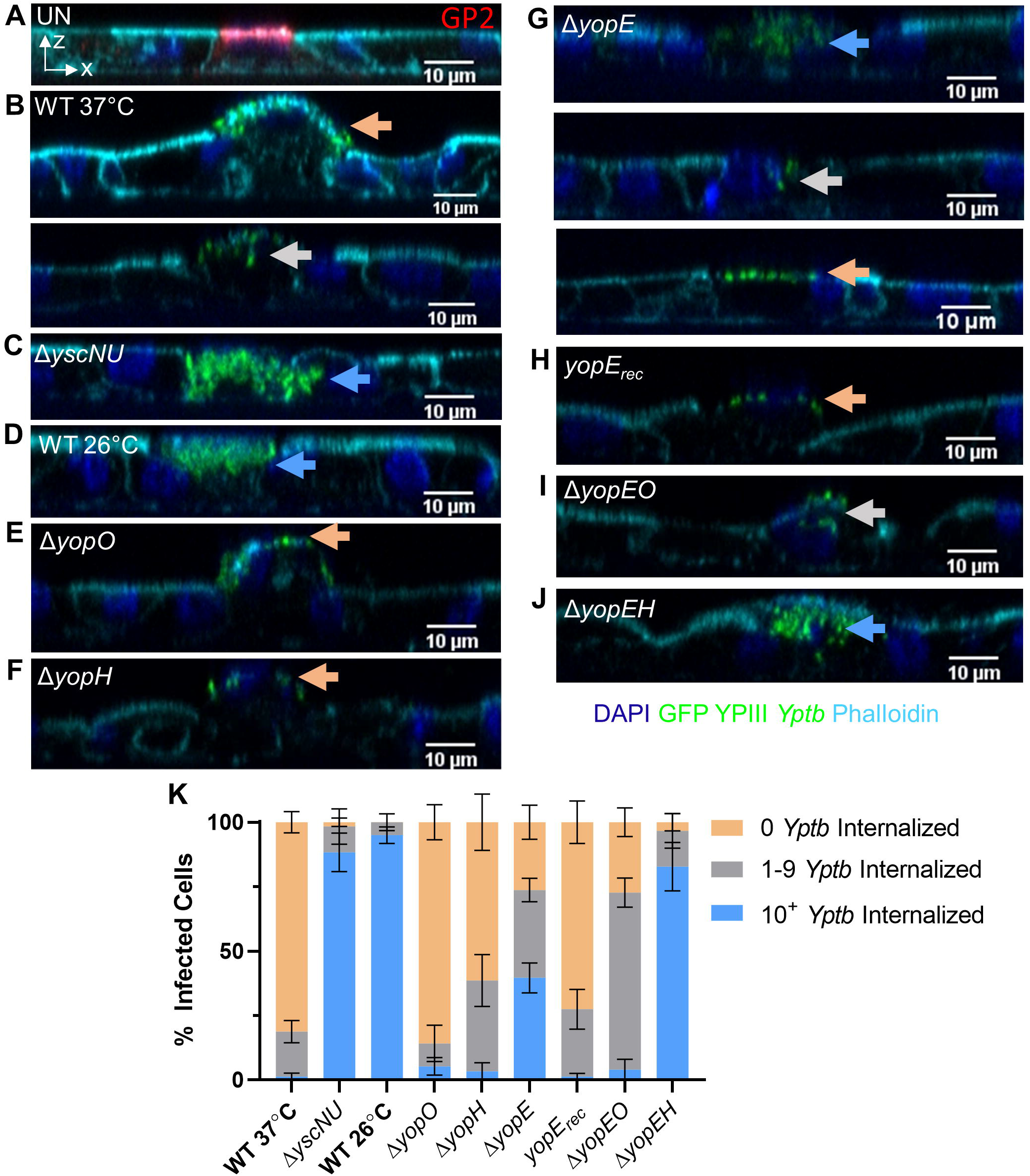
YopE and YopH contribute to inhibition of *Yptb* internalization by M cells. (A-J) Differentiated HIE25 RT+ ileal monolayers were (A) uninfected or infected for 5 hours with 5×10^6^ CFU of (B) WT 37°C, (C) *ΔyscNU*, (D) WT 26°C, (E) *ΔyopO,* (F) *ΔyopH,* (G) *ΔyopE*, (H) *ΔyopE* + *yopE_rec_,* (I) *ΔyopEO,* and (J) *ΔyopEH* YPIII *Yptb* expressing GFP (green). Monolayers were stained with DAPI (nuclei-blue) and phalloidin (F-actin-cyan). (A) was stained with anti-GP2 antibody (M cells-red). Orthogonal XZ planes were analyzed for the presence of *Yptb* on an M cell surface (0 *Yptb* internalized-orange arrows), partially internalized in an M cell (1-9 *Yptb* internalized-gray arrows), or fully internalized in an M cell (10^+^ *Yptb* internalized-blue arrows). (K) Number of *Yptb* internalized in each M cell was determined and the percentage of cells with the specified number of internalized *Yptb* per field was plotted. Error bars indicate SEM. Data were pooled from 3+ independent experiments with 2-4 fields analyzed per Transwell and averaged. Statistics were performed using a two-way ANOVA and Tukey’s post hoc multiple comparison tests and are shown in Table S1.

### YopE and YopH contribute to the inhibition of *Yptb* internalization by M cells

Given the well-established role of the T3SS effectors, YopE, YopH, and YopO in preventing *Yptb* uptake in a variety of cells ^27–31^, we next probed for their involvement in *Yptb* internalization by M cells. Similar to WT 37°C, the interaction of Δ*yopO* or Δ*yopH* with M cells was mostly restricted to cell surface binding, although a sizable, but not statistically different, minority of M cells had 1-9 Δ*yopH* internalized (Fig 4E, F, K, Table S1). After infection with Δ*yopE*, the three internalization classes were almost evenly divided (Fig 4G, K). This intermediary phenotype of Δ*yopE* was reflected in the percentage of M cells with 10^+^ *Yptb* internalized, which was significantly increased compared to WT 37°C, but significantly decreased compared to T3SS non-expressing strains (Table S1). Complementation of Δ*yopE* by expressing YopE from its native locus (*yopE_rec_*) restored the phenotype back to WT 37°C (Fig 4H, K, Table S1). Δ*yopEO* had a phenotype between Δ*yopE* and Δ*yopO,* with the majority of M cells taking up 1-9 *Yptb* (Fig 4I, K, Table S1). Finally, the percentage of M cells with 10^+^ *Yptb* internalized of Δ*yopEH* was similar to the T3SS non-expressing strains but significantly different from Δ*yopE* (Fig 4J, K, Table S1). Combined, these data indicate that the inhibition of internalization seen in Δ*yopE* is abrogated by Δ*yopEH* suggesting that together YopE and YopH contribute to inhibition of internalization by M cells while YopO has less of an effect.

### *Yptb* T3SS causes M cell extrusion from ileal monolayers independent of cell death

We noted that infected M cells bound by WT 37°C were extruding apically from the monolayers (Figs 4B, 5A). By contrast, uninfected M cells and M cells infected with Δ*yscNU* or WT 26°C remained flat within the monolayer indicating that extrusion is dependent on the T3SS (Figs 4A, C, D, 5A). The slight but non-significant increase in M cell extrusion after infection with WT 26°C compared to Δ*yscNU* is likely explained by 37°C-induced expression of the T3SS in WT 26°C that remained apically present during infection.

**Figure 5.**
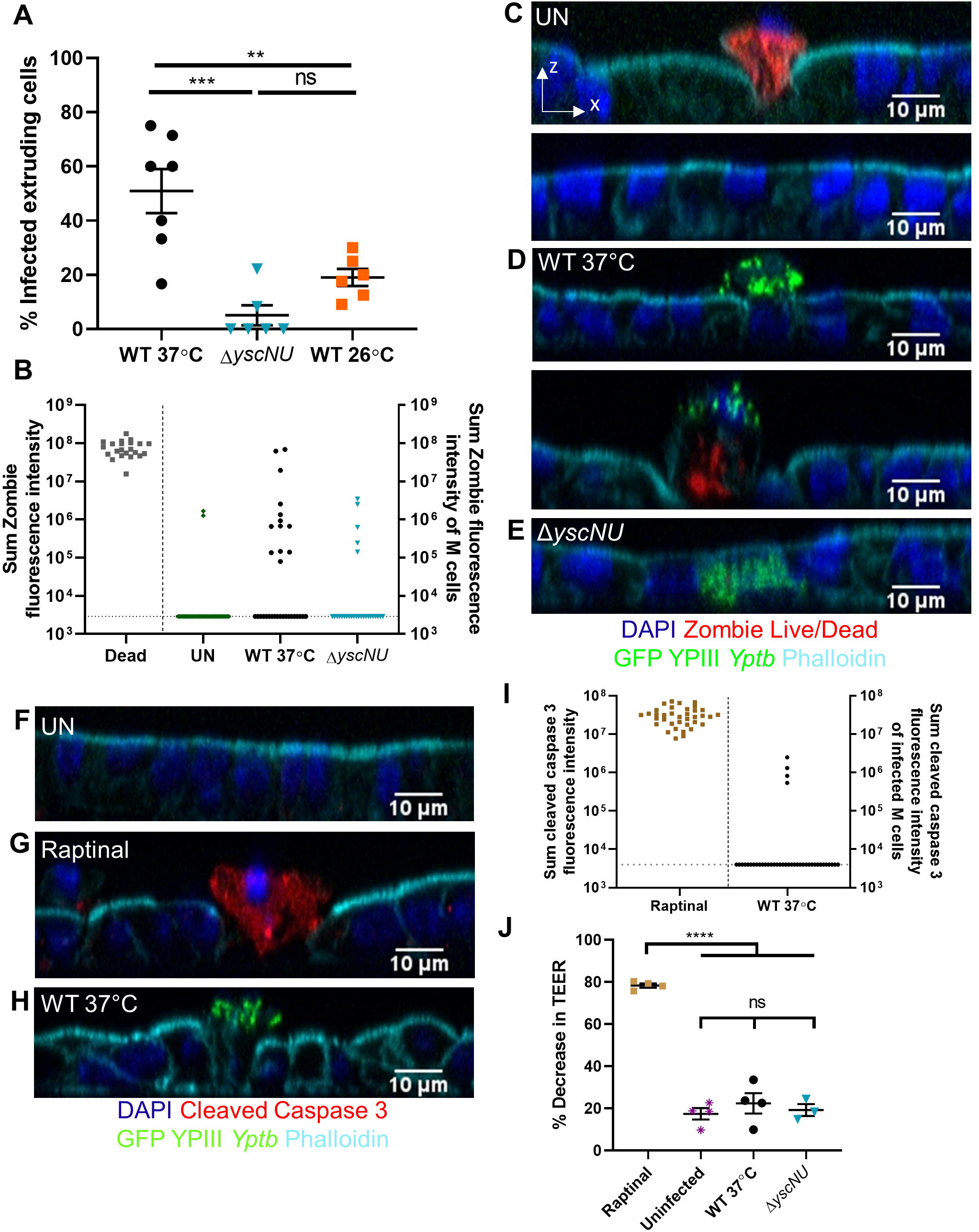
*Yptb* T3SS causes M cell extrusion independent of cell death. (A) Differentiated HIE25 RT+ ileal monolayers were infected for 5 hours with 5×10^6^ CFU of WT 37°C, *ΔyscNU*, or WT 26°C YPIII *Yptb* expressing GFP. The percentage of extruding infected M cells per field was plotted. Each point represents one Transwell. Bars indicate mean and SEM. Data were pooled from 3+ independent experiments with 2-4 fields analyzed per Transwell and averaged. Statistics were performed with a one-way ANOVA and Tukey’s post hoc multiple comparison tests. (B-E) Differentiated HIE25 RT+ ileal monolayers were (C) uninfected or infected for 5 hours with 5×10^6^ CFU (D) WT 37°C, or (E) Δ*yscNU* YPIII *Yptb* expressing GFP (green) and stained with Zombie Live/Dead stain (red), DAPI (nuclei-blue), and phalloidin (F-actin-cyan). Orthogonal XZ planes are shown. (B) Sum fluorescence intensity of Zombie. Each point represents a dead extruding cell or an M cell that was uninfected (GP2^+^) or infected with WT 37°C or Δ*yscNU* YPIII *Yptb* expressing GFP. 37 uninfected, 32 WT 37°C-infected, and 27 Δ*yscNU-*infected M cells were analyzed. Horizontal line indicates limit of detection. Data were pooled from 3 independent experiments. (F-H) Differentiated HIE25 RT+ ileal monolayers were (F) uninfected, (G) treated with 13 µM raptinal, or (H) infected for 5 hours with 5×10^6^ CFU WT 37°C YPIII *Yptb* expressing GFP (green) and stained for cleaved caspase 3 (red), DAPI (nuclei-blue), and phalloidin (F-actin-cyan). (I) Sum fluorescence intensity of cleaved caspase 3. Each point represents either a raptinal-induced apoptotic epithelial cell or a WT 37°C-infected M cell. 40 WT 37°C-infected M cells were analyzed. Horizontal line indicates limit of detection. (J) TEER was measured before and after infection and plotted as % decrease. Statistics were performed with a one-way ANOVA and Tukey’s post hoc multiple comparison tests. (I-J) Data were pooled from 3-4 independent experiments.

Cell extrusion can occur when epithelial cells die, which can be induced by infection with some enteric pathogens such as *Salmonella* and EHEC.^54,55^ To determine if *Yptb* causes infected M cells to die, a Zombie Live/Dead discrimination stain was used to observe if infected M cells lose membrane integrity during the 5-hour infection (Fig 5B-E). Dead extruding cells, defined by bright condensed nuclei, were sparsely located in uninfected monolayers and were used as a positive control for Zombie staining (red) (Fig 5B, C, Table S2). Uninfected monolayers had 0.4-3.1% of non-extruding cells stain with Zombie at two orders of magnitude below the positive controls (Table S2). 10 of 32 WT37°C-infected and 5 of 27 Δ*yscNU-*infected M cells (31% and 18.5%, respectively) had Zombie fluorescence that was also within the fluorescence range of uninfected non-extruding positive cells. For comparison, 2 of 37 of uninfected GP2^+^ M cells (5.4%) had Zombie fluorescence within this range (Fig 5B). The majority of WT 37°C-infected and Δ*yscNU-*infected M cells (60% and 81.5%, respectively) were not Zombie positive (Fig 5B, D, E, Table S2). Three WT 37°C-infected M cells had Zombie fluorescence levels similar to dead cells (Fig 5D, panel 2). These M cells appeared to be almost fully extruded, although not all nearly extruded M cells had Zombie intensity at the level of dead cells (Fig 5D, panel 2). Since 60% of WT 37°C-infected M cells did not lose detectable membrane integrity, these data indicate that *Yptb*-induced M cell extrusion begins prior to loss of membrane integrity and suggest that *Yptb* is not triggering lytic cell death prior to extrusion.

Cells undergoing early apoptosis do not lose membrane integrity yet apoptotic signals often lead to cell extrusion.^56^ Cleavage of caspase 3, an executioner caspase, leads to downstream pathways that initiate apoptotic features.^57,58^ To evaluate whether WT 37°C causes M cells to undergo apoptosis, monolayers were stained for cleaved caspase 3. Uninfected monolayers had 1.5-5.1% of cells stain positively for cleaved caspase 3 (Fig 5F, Table S3). Monolayers treated for 5 hours with raptinal, an apoptosis inducer, had many cells stain positively for cleaved caspase 3 (red) (Fig 5G). By contrast, 36 of 40 WT 37°C-infected extruding M cells (90%) were negative for cleaved caspase 3 (Fig 5H-I). The 4 infected M cells above the limit of detection had fluorescence levels below that of raptinal-induced apoptotic cells (Fig 5I). TEER measurements taken before and after raptinal treatment or infection revealed a severe loss in monolayer integrity after raptinal treatment, but a small non-significant decrease in TEER in infection conditions compared to uninfected monolayers (Fig 5J). Collectively, the Zombie, cleaved caspase 3, and TEER results support the idea that *Yptb*-induced M cell extrusion occurs independently of cell death and does not cause significant disruptions to monolayer integrity.

### YopE RhoA-GAP activity is primarily responsible for M cell extrusion

Since M cells, but not adjacent cells, were extruding from the monolayer, this event was likely due to *Yptb* binding or injection of Yops into M cells. To investigate the involvement of Yops in M cell extrusion, orthogonal planes of monolayers infected with *yop* mutants were analyzed. Infection with Δ*yopO* or Δ*yopH* caused levels of M cell extrusion similar to WT 37°C (Fig 6A-C, I). By contrast, M cells infected with Δ*yopE* rarely extruded (Fig 6D, I).

**Figure 6.**
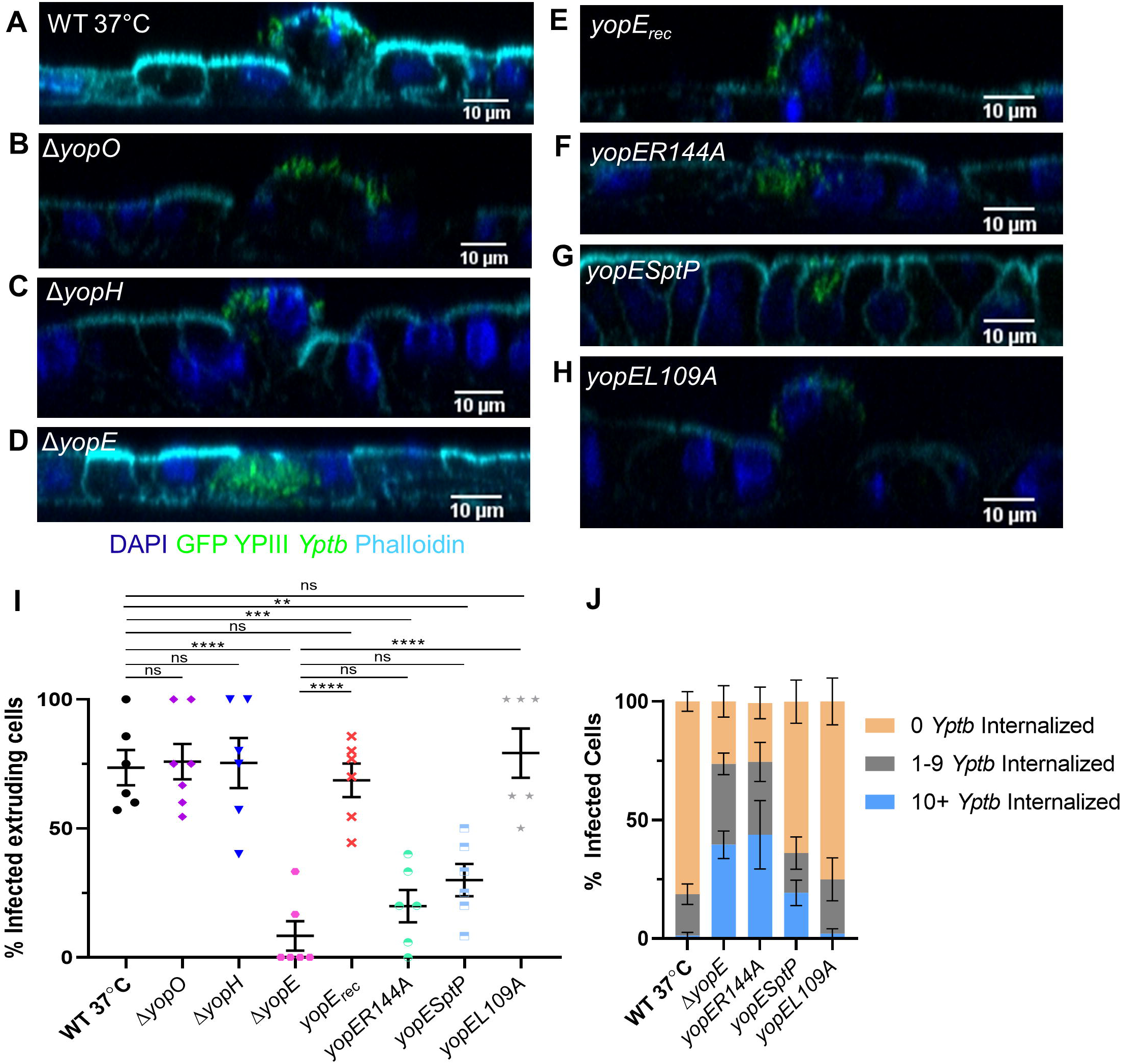
YopE causes M cell extrusion. (A-H) Differentiated HIE25 RT+ ileal monolayers were infected for 5 hours with 5×10^6^ CFU of (A) WT 37°C, (B) *ΔyopO,* (C) *ΔyopH,* (D) *ΔyopE*, (E) *ΔyopE*+*yopE_rec_,* (F) *ΔyopE+yopER144A,* (G) *ΔyopE+YopE_1-100_SptP_166-293_* or (H) *ΔyopE+yopEL109A* YPIII *Yptb* expressing GFP (green). Monolayers were stained with DAPI (nuclei-blue) and phalloidin (F-actin-cyan). Orthogonal XZ planes were analyzed for M cell extrusion. (I) The percentage of extruding infected M cells per field was plotted. Each point represents one Transwell. Bars indicate mean and SEM. Statistics were performed with a one-way ANOVA and Tukey’s post hoc multiple comparison tests. (J) The number of *Yptb* internalized in each M cell was determined and the percentage of cells with specified number of internalized *Yptb* per field was plotted. Error bars indicate SEM. The data for WT 37°C and Δ*yopE* are the same as in Figure 4K. Statistics were performed using a two-way ANOVA and Tukey’s post hoc multiple comparison tests and are shown in Table S4. (I-J) Data were pooled from 3+ independent experiments with 2-4 fields analyzed per Transwell and averaged.

Complementation of Δ*yopE* (*yopE_rec_*) restored M cell extrusion back to the level seen with WT 37°C (Fig 6A, E, I). Since over 30% of M cells infected with Δ*yopE* had 10^+^ *Yptb* internalized (Fig 4K), this raised the possibility that these Δ*yopE*-infected M cells simply might not extrude because rapid uptake of *Yptb* results in no Yop injection. However, of the M cells with 0 Δ*yopE* internalized, only 11% were extruding, implying that YopE plays an active role in causing extrusion.

YopE targets Rac1 and RhoA of the Rho GTPase family of proteins that are involved in actin organization and assembly at cell-cell junctions.^59^ To determine if YopE-GAP activity is needed for M cell extrusion, monolayers were infected with the YopE catalytic mutant, *yopER144A*, which binds to small GTPases but lacks GAP activity.^32^ The catalytic mutant *yopER144A* behaved similarly to Δ*yopE,* with a significant reduction in M cell extrusion compared to WT 37°C (Fig 6F, I). Analysis of *Yptb* internalization revealed that *yopER144A* also behaved similarly to Δ*yopE* having an intermediary phenotype between the three internalization classes (Fig 6J, Table S4). The contribution of YopE-GAP activity to M cell extrusion was evaluated using two additional YopE mutants, *yopESptP* or *yopEL109A*. YopE_1-100_SptP_166-293_ is a fusion protein that has the secretion and translocation domains of YopE and the GAP activity of *Salmonella* SptP, which lacks RhoA-GAP activity and has slightly weaker Rac1-GAP activity (about 80% of WT YopE).^33^ *Yptb* expressing *yopESptP* was significantly impaired in causing M cell extrusion and remained mostly bound to the apical surface of M cells (Fig 6G, I, J, Table S4). By contrast, infection with *yopEL109A*, which has partially reduced RhoA-GAP activity (about 70% of WT YopE) and normal Rac1-GAP activity ^33^, caused M cell extrusion similar to WT 37°C and had a *Yptb* internalization phenotype similar to WT 37°C (Fig 6H-J, Table S4). Combined, these data suggest that the RhoA-GAP activity of YopE is primarily responsible for causing M cells to extrude from the epithelium. Furthermore, since both *yopESptP* and *yopEL109A* retain Rac1-GAP activity, while *yopER144A* does not, these data also support the idea that the Rac1-GAP activity of YopE contributes to inhibiting internalization by M cells.

### M cells infected with WT 26°C have a reduced ability to be reinfected by WT 37°C

Since *Yptb* infections are often acquired by eating contaminated food stored at cold temperatures, the bacteria may reach the intestine before fully expressing the T3SS, with subsequent growth in the intestine causing induction of T3SS expression. To model this scenario, we evaluated the interaction of *Yptb* with M cells that were first infected with GFP-expressing WT 26°C and then challenged 2.5 hours later with mCherry-expressing WT 26°C or WT 37°C (Fig 7). The number of mCherry-expressing *Yptb* associated with M cells grown under either condition were compared to the number of mCherry-expressing WT 37°C associated with M cells without prior *Yptb* infection. Prior infection with WT 26°C dramatically decreased the ability of WT 37°C to associate with M cells during subsequent infection (7A, C, D). By contrast, mCherry-expressing WT 26°C associated with M cells after prior infection as efficiently as infection with WT 37°C without prior infection (Fig 7A, B, D). These results are consistent with the idea that prior infection with WT 26°C reduces the number of apically exposed β1 integrins so WT 37°C that express reduced invasin ^60,61^ are at a competitive disadvantage in associating with M cells.

**Figure 7.**
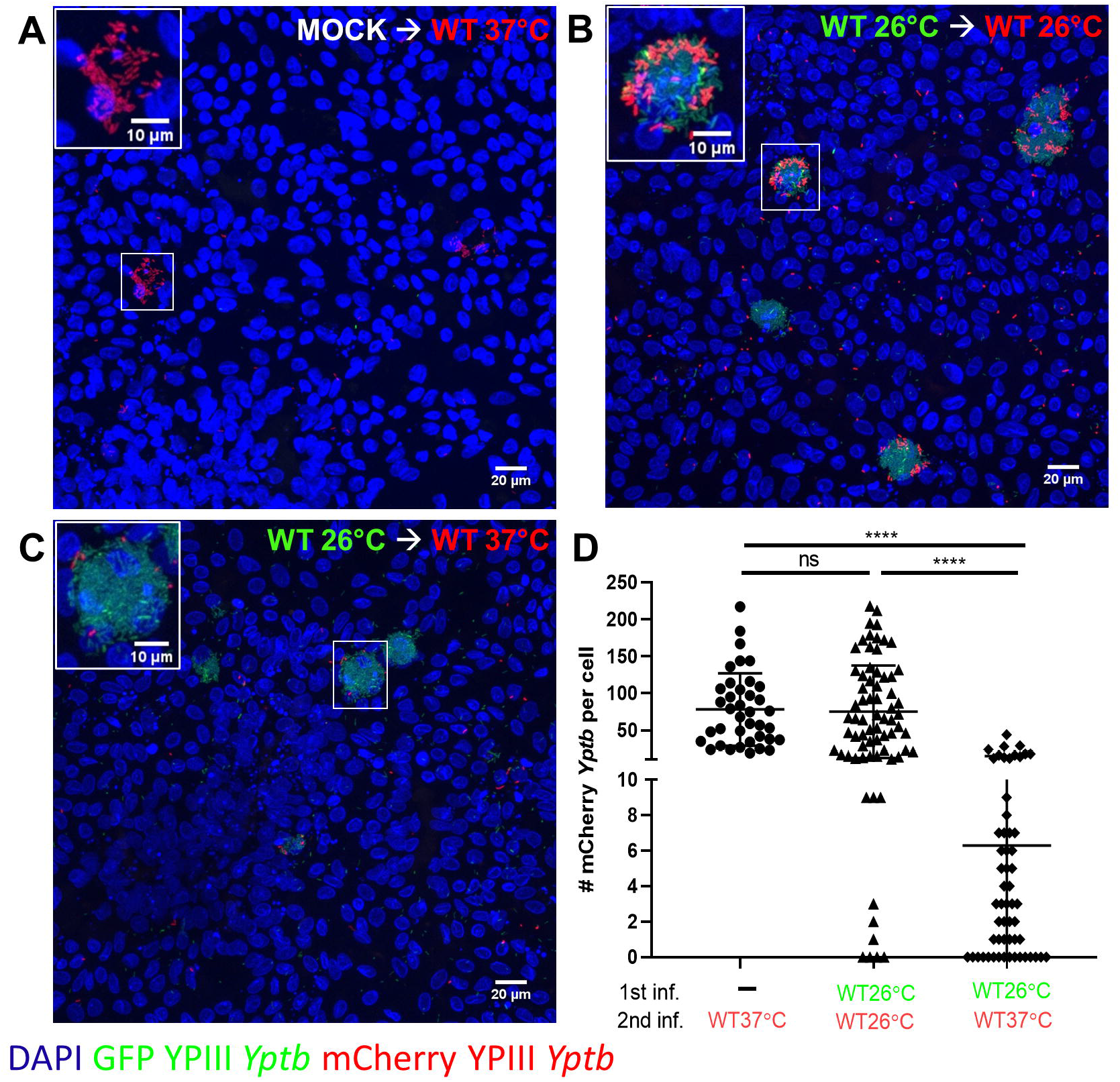
M cells infected with WT 26°C have reduced ability to be reinfected by WT 37°C. (A-D) Differentiated HIE25 RT+ ileal monolayers were (A) mock infected with non-infectious media, washed, and infected for 2.5 hours with 5×10^6^ CFU of WT 37°C expressing mCherry (red), or (B) infected for 2.5 hours with 5×10^6^ CFU of WT 26°C expressing GFP (green), washed, and infected for 2.5 hours with 5×10^6^ CFU of WT 26°C expressing mCherry (red), or (C) infected for 2.5 hours with 5×10^6^ CFU of WT 26°C expressing GFP (green), washed, and infected for 2.5 hours with 5×10^6^ CFU of WT 37°C expressing mCherry (red). Monolayers were stained with DAPI (nuclei-blue). XY planes are maximum intensity projections. Magnified insets of an M cell are shown in upper left corner. (D) The number of mCherry *Yptb* per cell were counted using Volocity software. Each point represents an infected M cell. Bars indicate mean and SD. Data were pooled from 3 independent experiments with 3 fields analyzed per Transwell and averaged. Statistics were performed with a Kruskal-Wallis multiple comparisons test.

## Discussion

M cells are used by many pathogens to penetrate the intestinal epithelial barrier, yet the intricate host-pathogen interactions that occur at the M cell interface have been challenging to study *in vitro* due to biological limitations of current transformed cell culture models.^41^ By employing human enteroid-derived monolayers containing functional M cells interspersed among an epithelium containing at least 4 other intestinal cell types, we reproduced the findings in ligated loop models that *Yptb* binds and congregates specifically to M cells in an invasin dependent manner. ^2,14^ We further exploit this model to discover how Yops impact M cell function and fate, to find that the consequences of these interactions could affect the ability of *Yptb* to establish a successful foothold in underlying tissues.

By exploiting the Transwell model, we distinguish the impact of early infectious state (low levels of Yop-expression) from host-adapted state on M cells to understand the conundrum of how *Yptb* gets through M cells when YopE and YopH prevent internalization and phagocytosis, yet are essential for survival in the intestinal tract and PP.^15,28,30,31,39,40,62,63^ Yops were injected into and were effective in M cells because WT 37°C caused M cell extrusion in a YopE GAP-dependent manner and were rarely internalized by M cells due to YopE and YopH. However, an important consideration in envisioning initial steps in the establishment of *Yptb* infection is that *Yptb* consumed in contaminated food at cold temperatures may reach the intestine and encounter M cells before fully expressing the T3SS. Here, transcytosis of *Yptb* grown at ambient temperatures mimicking early infectious stage (WT 26°C) occurred at levels 10 times that of host-adapted (WT 37°C) expressing the T3SS. Even though transcytosis of WT 37°C was less frequent, it was still significantly detected in the presence of M cells compared to the absence, demonstrating that M cell-containing monolayers are required for detectable transcytosis in this model. Murine studies with T3SS mutants, such as *yscNU*, have shown that these bacteria are able to colonize PP but are far less likely to survive at 24 hours post-infection.^39,63^ Since the T3SS gives *Yptb* an advantage upon encountering immune cells ^22^, it is possible that the rare T3SS-expressing *Yptb* that breach M cells are ultimately more successful in animal infection. Alternatively, the successful bacteria may be those that are expressing low levels of Yops during encounter with M cells but are ramping up T3SS expression as they travel through the M cell. In fact, during infection *Yersinia* can respond rapidly to environmental cues, such as host cell contact, to increase the copy number of the plasmid carrying the T3SS which is essential for virulence consistent with this idea.^64^

Intestinal cell extrusion is a normal part of epithelial regeneration that occurs when cells are shed from the epithelium and undergo anoikis, a detachment induced form of apoptosis.^65^ In addition, pathogens can also induce different inflammatory and non-inflammatory cell extrusion processes, which are often characterized by cell death.^66^ *Salmonella* promotes destruction and death of M cells in murine ileal ligated loops resulting in open gaps in the FAE that allow for additional invasion of bacteria.^67^ *Salmonella* has also been shown to induce death and extrusion of intestinal epithelial cells characterized by inflammatory caspase activation.^54,68^ By contrast, Enterohaemorrhagic *E. coli* infection of MDCK cells activates RhoA to cause a non-cell death extrusion.^55^ Here we show that *Yptb*-induced M cell extrusion was dependent on the RhoA-GAP activity of YopE and was not triggered by cell death.

How does the loss of RhoA activity promote M cell extrusion? RhoA stimulates contractility and promotes actin stress fiber formation by regulating actomyosin complexes that control tension and forces between cells, which helps to maintain epithelial monolayer integrity.^59,69,70^ Widespread inactivation of RhoA, using the Rho specific inhibitor C3 transferase, prevents apical extrusion of apoptotic cells.^71^ During *Yptb* infection, targeted injection of Yops into only M cells limits the inhibitory effects of YopE on RhoA to within infected M cells. Therefore, localized M cell extrusion may be similar to a cell non-autonomous extrusion process whereby neighboring cells actively participate by recognizing local changes in tension.^72^ Two recent reports have shown that contractile tension that occurs in response to a perturbation, such as apoptosis or inducible expression of the epithelial morphogenesis transcription factor Snail^6SA^ in some cells, activates RhoA in neighboring cells resulting in apical extrusion of affected cells.^73,74^ Based on these findings, it is plausible that YopE inhibition of RhoA activity in M cells results in the generation of contractile tension that is sensed by neighboring cells that respond by activating RhoA to contract and squeeze the infected M cell out of the monolayer. The idea that neighboring cells respond to infected M cells is supported by our findings that the TEER did not decrease after infection and that monolayers stained for F-actin appeared to be closing up beneath extruding M cells (unpublished data). Therefore, unlike *Salmonella* induced destruction of M cells ^67^, it is not likely that *Yptb-*induced M cell extrusion creates holes in the FAE as additional access points for *Yptb*.

In mice, sequential infection with green-fluorescing *Y. enterocolitica* followed by infection with red-fluorescing *Y. enterocolitica* 2 days later resulted in infection by red bacteria of PP that were not already colonized by green bacteria.^17^ In addition, enteric *Yersinia* form clonal microcolonies in PP and deeper tissues.^15,17,18^ These findings show that the host may limit infection at the level of PP invasion and/or survival in PP ^17^, but it remains unclear. Our observation that M cells infected with WT 37°C induced M cells to extrude indicate a possible means by which a secondary infection could be limited via temporary depletion of M cells above a PP. M cell depletion has been described after reovirus type I infection in mice and porcine epidemic diarrhea virus (PEDV) infection in pigs ^75,76^, although the mechanisms are not defined. It remains to be observed what the long-term (48-hour post infection) effects of infection with WT 26°C may be and whether this too could impair M cell function.

Our studies of *Yptb* interactions with M cells provide two important insights into M cell development and function. First, as M cells develop after exposure to RANKL and TNFα ^46,47^, they gain expression of various markers including GP2, a late stage marker of M cells indicative of maturity.^50,51^ When expression of apical β1 integrin occurs in the developmental timeline is unknown; however, the observation that a population of GP2 low/negative cells bound *Yptb* and that these cells were β1 integrin^+^ supports the idea that apical β1 integrin may be expressed earlier in M cell development. While GP2 low/negative cells have reduced uptake capabilities ^77^, *Yptb* uptake by these M cells is likely efficient given the nature of the strong invasin-β1 integrin interaction.^78,79^ These M cells could also be at risk for depletion from the FAE by YopE-induced extrusion implying that multiple developmental stages of M cells may be affected by *Yptb*. Second, M cell functionality can be induced by molecules other than those that contribute to M cell lineage commitment including CD137L, S100A4, and retinoic acid/lymphotoxin (LT-α2β1).^52,80,81^ The combination of CD137L with RANKL and TNFα during differentiation yielded no further increase in transcytosis of *Yptb*. Since CD137L is a TNFα family member, it is possible that CD137L and TNFα stimulate similar signaling pathways to invoke M cell functional maturation and their presence together is redundant.

In summary, this study emphasizes the power of using enteroid models to uncover new insights into pathogen-epithelium interactions. Using this approach, we discovered that *Yptb* Yops impair M cell function by reducing *Yptb* uptake and causing M cell extrusion, both of which may benefit the host. These findings support the idea that efficient transit of *Yptb* through M cells occurs before high levels of T3SS expression. Future work is focused on increasing the complexity of this enteroid system by including immune cells to explore how the status of T3SS expression of WT *Yptb* affects *Yptb* survival as they breach the FAE and encounter immune cells.

## Materials and Methods

### Seeding and Differentiation of Enteroids as Monolayers on Transwells

Growth and propagation of enteroids is described in.^82^ All media compositions can be found in.^49^ Noggin conditioned media was prepared according to protocols in.^83^ Enteroids were grown for about 1 week in Matrigel (Corning) with the following modifications. Growth media was sometimes used at 75% v/v or 50% v/v Wnt3a conditioned media. Transwells of 0.4 µm, 1 µm, or 3 µm pore size were seeded and maintained as in.^49^ To differentiate monolayers on Transwells, growth media was removed from the apical and basolateral chambers after 1-3 days post-seeding, (once monolayers were 80-95% confluent) and was replaced with differentiation media. To induce M cell development, the basolateral chamber differentiation media was supplemented with 200 ng/mL RANKL (Peprotech; cat #: 310-01) and 50 ng/mL TNFα (Peprotech; cat #: 315-01A). The media was replaced every other day and monolayers were used for infection experiments on day 4 - 7 post-seeding based on monolayer confluence and increased transepithelial electrical resistance (TEER) measured with a Millicell ERS-2 Voltohmmeter device.

### Bacterial Strains and Infections

*Yptb* YPIII strains used for this work are listed in Table S5. The following strains were generated using bacterial conjugation described in.^84^ To generate Δ*yopH* and Δ*yopO*, pCVD442-yopHKO or pCVD442-yopOKO were conjugated into WT YPIII (JM301), respectively. To generate Δ*yopEH* and Δ*yopEO*, pCVD442-yopHKO or pCVD442-yopOKO were conjugated into Δ*yopE* YPIII (WS125c), respectively. Plasmids are described in.^39^ To generate Δ*yadA* and Δ*inv*, pCVD442-yadAKO or pCVD442-invKO was conjugated into WT YPIII (JM301), respectively. To generate Δ*inv/yadA*, pCVD442-yadAKO was conjugated into Δ*inv* (ERG9). Plasmids are described in.^85^

*Yptb* strains constitutively expressed either green fluorescent protein (GFP) via transformation of the pACYC184-GFP plasmid (chloramphenicol resistant) or the red fluorescent protein, mCherry, via transformation of the pMMB67EH-mCherry plasmid (carbenicillin resistant). *Yptb* strains were grown in 2xYT broth with appropriate drugs (20 µg/mL chloramphenicol, 100 µg/mL carbenicillin) at 26°C with aeration overnight. In the morning, cultures were back-diluted at 1:40 in 2xYT with appropriate drugs in either low calcium (Figs 1-2) or high calcium conditions (Figs 3-7). Low calcium media contained 2xYT supplemented with 20 mM sodium oxalate and 20 mM MgCl_2_. High calcium media contained 2xYT supplemented with 5 mM CaCl_2_. Cultures were grown at 26°C for 2 hours with aeration and were shifted to 37°C for 2 hours with aeration to induce T3SS expression. All mutant strains were shifted to 37°C unless otherwise specified. The bacterial cultures were resuspended in differentiation media at 5×10^6^ CFU in 200 µL. Transwells were infected apically for 2.5, 3 or 5 hours as indicated in a tissue culture incubator at 37°C and 5% CO_2_. After infection, the media was gently removed from the apical chamber and Transwells washed 2x with room temperature PBS. Monolayers were fixed with room temperature 4% paraformaldehyde (PFA) for 25 minutes followed by 3 washes with PBS and stored in PBS at 4°C until they were processed for immunofluorescence microscopy.

### Transcytosis Assays

On the day of infection, the TEER of monolayers that were grown on 3 µm Transwells was measured to ensure the monolayer was polarized (TEER > 400 Ω·cm ^2^). Infections were carried out for 3 hours. After infection, Transwells were removed from the 24 well plate and placed in a new 24 well plate for fixation and the basolateral chamber media was plated on L irgasan (0.5 µg/mL) agar plates to determine the CFU. After fixation, the monolayers were permeabilized with 0.1% TritonX-100 in 1% bovine serum albumin (BSA) and stained with Alexa Fluor 647 phalloidin and DAPI. Every monolayer was scrutinized on a Nikon A1R confocal microscope for the presence of holes, defined as regions with loss of DAPI and phalloidin stains with GFP bacteria bound to the Transwell filter membrane. Monolayers with holes were omitted from further analysis of basolateral chamber CFU.

### Zombie Live/Dead Cell Death Assays

The Zombie Red Live/Dead Discrimination Stain was used according to the manufacturer’s instructions with the following modifications. At 4.5 hours after infection, the media in the apical chamber was gently removed and monolayers were washed 1x with PBS. Zombie Red Live/Dead Discrimination Stain was diluted 1:200 in PBS and 100 µL was added to monolayers and incubated at 37°C in tissue incubator for 30 minutes. Following incubation, the monolayers were washed 2x with 1% BSA in PBS and were fixed and stored as described above.

### Staining and Confocal Immunofluorescence Microscopy

For surface staining of β1 integrin and GP2, monolayers were not permeabilized before primary staining. For staining of cleaved caspase 3, monolayers were permeabilized with 0.1% Triton X-100 in 1% BSA in PBS for 5 minutes before primary staining. After permeabilization, Transwells were blocked in 5% BSA in PBS for 30 minutes at room temperature. Primary antibodies were diluted in 1% BSA with in PBS at 1:100 with 100 µL applied per Transwell. Primary antibodies included GP2 (MBL International; cat #: D277 -3), β1 integrin/CD29 (Fisher Scientific; cat #: BDB555442), and cleaved caspase 3 (Cell Signaling; cat #: 9661). The primary stain was carried out at room temperature in the dark for 1 hour. Transwells were then washed 3 times with PBS. Corresponding anti-mouse secondary Alexa Fluor antibodies were diluted 1:200 in 1% BSA and 0.1% Triton X-100 in PBS, along with 1 µg/mL 4′,6-diamidino-2-phenylindole (DAPI) (Invitrogen; cat#: 62248) to stain the nuclei and Alexa Fluor 647 phalloidin at 1:100 (Invitrogen; cat #: A22287) to stain F-actin. The secondary stain was carried out at room temperature in the dark for 30 minutes and then the Transwells were washed 3 times. Stained monolayers were carefully cut from the Transwell using a scalpel and placed on glass slides with the cell layer facing up. Monolayers were covered with Prolong Gold antifade (Invitrogen; cat #: P36930) and mounted with a glass coverslip. Glass slides were placed at 4°C in the dark until they were brought to the microscope for image acquisition. Images were acquired with a Nikon A1R confocal microscope (Nikon Instruments Inc.) using a 40x oil immersion objective with 0.5 µm z-stacks. Images were analyzed using FIJI ImageJ or Volocity 6.3 software (Quorum Technologies).

### Image Acquisition and Analysis

In initial experiments, M cells were located by presence of apical β1 integrin or GP2 proteins. After analysis of *Yptb* binding experiments, M cells were defined as cells that contained 6+ *Yptb* bound to the apical surface so that other antibody stains could be used. Images taken for RT+ monolayers were obtained in blinded manner such that the only requirement for selecting a field for image capture was that at least 1 M cell was present in the field. While scanning the monolayers, the first three fields with M cells present were selected for image capture in each Transwell. Only Transwells with 4+ M cells across the 3 images were used for subsequent analysis. The numbers of M cells per field depended on the pore size and experiment and ranged from 1-10 M cells per field. In RT- monolayers, three non-contiguous fields were randomly selected for analysis.

Images were blinded for all subsequent analysis and quantification. To analyze *Yptb* binding to differentiated cells of the intestinal monolayer, cells were manually counted for the presence of *Yptb* on the apical surface. When a single bacterium was present at a junction point between two cells, this was counted as one bacterium bound to one cell. To quantify M cell extrusion and *Yptb* internalization into M cells, the orthogonal views of the monolayers were observed with an image processing software. M cells were manually counted as apically extruding if any part of the M cell volume resided above the apical plane of the neighboring cells. Fully extruded cells that contained *Yptb* were eliminated from the analysis to prevent counting cells that had extruded in normal epithelial turnover prior to infection. Internalized *Yptb* were manually counted by locating the boundaries of a cell with F-actin. GFP *Yptb* that fell within the boundaries of the F-actin stain were counted as internalized. The fluorescence intensity for phalloidin was measured as described in.^49^

For Zombie Live/Dead cell analysis, Volocity was used to measure the sum fluorescence intensity of the Zombie Red channel. Dead cells were determined by locating cells that were Zombie positive, extruding, and had a ratio of sum DAPI fluorescence/DAPI volume that was greater than that for live cells to signify bright condensing nuclei. For cleaved caspase 3 analysis, Volocity was used to measure the sum fluorescence intensity of the Alexa Fluor 594 channel (cleaved caspase 3). Cells were induced to undergo apoptosis using 13 µM raptinal (AdipoGen; cat#: AG-CR1-2902) and apoptotic cells were defined as cells positive for cleaved caspase 3.

### Statistical Analysis

Statistical analyses were completed using GraphPad Prism Software version 8. Statistical tests used are described in individual figure legends, where * p<0.05, ** p<0.01, *** p<0.001, **** p<0.0001, and ns = no significance for all figures and supplemental tables.

### Tissue Collection // Ethics Statement

Human ileal enteroid cultures were obtained from de-identified biopsy tissue from healthy subjects who provided informed consent. Methods were performed following regulations that were approved by the Baylor College of Medicine’s Institutional Review Board and the Johns Hopkins University Institutional Review Board.

## Data Availability

The data that support the findings of this study are available from the corresponding author, JM, upon reasonable request.

## Acknowledgements

We thank Eric Stas (Boston Children’s Hospital), Igor Brodsky (University of Pennsylvania), Alenka Lovy, members of the Kaplan, Ng, and Ward Labs (Tufts Medical Center), Lamyaa Shaban, Giang Nguyen, Anne McCabe, Pathricia Leus, Rebecca Silver, Yoelkys Morales, and previous members of the Mecsas Lab for technical support, strains, and/or critical reading of the manuscript. This work was supported by NIAID U19AI131126 to RRI (Tufts University School of Medicine) and Dr. Kaplan (Tufts University) with JM serving as Project 2 Leader; NIAID R21AI128093 to JM; NIH U19AI116497 to MKE; the Integrated Physiology Core of the Hopkins Conte Digestive Disease Basic and Translational Research Core Center (NIH DK-809502) for HIE59; and the Tufts Center for Neuroscience Research, P30 NS047243 for confocal imaging. DTB was supported in part by the Harvard Digestive Disease Center P30 DK034854. ACF was supported in part by NIAID T32AI007077.

## Supplemental Information

**Supplemental Table 1**. Table of significances for Figure 4K.^a^

**Supplemental Table 2**. Additional details related to quantification of Zombie Live/Dead analysis

**Supplemental Table 3**. Additional details related to quantification of cleaved caspase 3 analysis

**Supplemental Table 4**. Table of significances for Figure 6J.^a^

**Supplemental Table 5**. *Yersinia* strains used in this study.

