## Supplemental Information for "*Yersinia pseudotuberculosis* YopE prevents uptake by M cells and instigates M cell extrusion in human ileal enteroid-derived monolayers"

Supplemental Table 1. Table of significances for Figure 4K.[a]

| | 0 *Yptb* Internalized | | | | | | | | |
|---|---|---|---|---|---|---|---|---|---|
| | WT 37°C | Δ*yscNU* | WT 26°C | Δ*yopO* | Δ*yopH* | Δ*yopE* | *yopE*rec | Δ*yopEO* | Δ*yopEH* |
| WT 37°C | | | | | | | | | |
| Δ*yscNU* | **** | | | | | | | | |
| WT 26°C | **** | ns | | | | | | | |
| Δ*yopO* | ns | **** | **** | | | | | | |
| Δ*yopH* | ns | **** | **** | ns | | | | | |
| Δ*yopE* | **** | ns | ns | **** | ** | | | | |
| *yopE*rec | ns | **** | **** | ns | ns | **** | | | |
| Δ*yopEO* | **** | ns | ns | **** | ** | ns | **** | | |
| Δ*yopEH* | **** | ns | ns | **** | **** | ns | **** | ns | |

| | 1-9 *Yptb* Internalized | | | | | | | | |
|---|---|---|---|---|---|---|---|---|---|
| | WT 37°C | Δ*yscNU* | WT 26°C | Δ*yopO* | Δ*yopH* | Δ*yopE* | *yopE*rec | Δ*yopEO* | Δ*yopEH* |
| WT 37°C | | | | | | | | | |
| Δ*yscNU* | ns | | | | | | | | |
| WT 26°C | ns | ns | | | | | | | |
| Δ*yopO* | ns | ns | ns | | | | | | |
| Δ*yopH* | ns | ns | * | * | | | | | |
| Δ*yopE* | ns | ns | * | ns | ns | | | | |
| *yopE*rec | ns | ns | ns | ns | ns | ns | | | |
| Δ*yopEO* | **** | **** | **** | **** | ** | ** | *** | | |
| Δ*yopEH* | ns | ns | ns | ns | ns | ns | ns | **** | |

| | 10+ *Yptb* Internalized | | | | | | | | |
|---|---|---|---|---|---|---|---|---|---|
| | WT 37°C | Δ*yscNU* | WT 26°C | Δ*yopO* | Δ*yopH* | Δ*yopE* | *yopE*rec | Δ*yopEO* | Δ*yopEH* |
| WT 37°C | | | | | | | | | |
| Δ*yscNU* | **** | | | | | | | | |
| WT 26°C | **** | ns | | | | | | | |
| Δ*yopO* | ns | **** | **** | | | | | | |
| Δ*yopH* | ns | **** | **** | ns | | | | | |
| Δ*yopE* | **** | **** | **** | ** | ** | | | | |
| *yopE*rec | ns | **** | **** | ns | ns | *** | | | |
| Δ*yopEO* | ns | **** | **** | ns | ns | ** | ns | | |
| Δ*yopEH* | **** | ns | ns | **** | **** | *** | **** | **** | |

[a] * $p < 0.05$; ** $p < 0.01$; *** $p < 0.001$; **** $p < 0.0001$; ns = no significance

Supplemental Table 2. Additional details related to quantification of Zombie Live/Dead analysis

| | Uninfected Monolayers | | |
|---|---|---|---|
| | Avg % of Zombie$^{+}$ Extruding dead cells | Range of mCherry intensity amongst Zombie$^{+}$ Extruding dead cells | Geometric Mean of mCherry Intensity amongst Zombie$^{+}$ Extruding dead cells |
| Expt 1 | 0.4% | $6x10^{6}$-$2x10^{8}$ | $7x10^{7}$ |
| Expt 2 | 0.4% | $6x10^{6}$-$1x10^{8}$ | $3x10^{7}$ |
| Expt 3 | 0.04% | $5x10^{6}$-$6x10^{7}$ | $2x10^{7}$ |

| | Uninfected Monolayers | | |
|---|---|---|---|
| | Avg % Zombie$^{+}$ Non-extruding Cells | Range of mCherry Intensity amongst Zombie$^{+}$ Non-extruding cells | Geometric Mean of mCherry Intensity amongst Zombie$^{+}$ Non-extruding cells |
| Expt 1 | 3.1% | $4x10^{4}$-$9x10^{7}$ | $5x10^{5}$ |
| Expt 2 | 1.0% | $9x10^{4}$-$9x10^{6}$ | $5x10^{5}$ |
| Expt 3 | 0.4% | $1x10^{5}$-$8x10^{5}$ | $4x10^{5}$ |

| | # M cells analyzed per experiment | | |
|---|---|---|---|
| | UN ([a,b]) | WT 37°C ([a,b]) | Δ*yscNU* ([a,b]) |
| Expt 1 | 29 (0,2) | 6 (1,2) | 6 (0,3) |
| Expt 2 | ND[c] | 18 (2,5) | 14 (0,2) |
| Expt 3 | 8 (0,0) | 8 (0,3) | 7 (0,0) |

[a] Number of Zombie+ cells in intensity range of Extruding Dead Cells
[b] Number of Zombie+ cells in intensity range of Non-extruding cells
[c] ND = Not determined

Supplemental Table 3. Additional details related to quantification of cleaved caspase 3 analysis

| | Uninfected Monolayers | | |
|---|---|---|---|
| | Avg % of $CC3^{+}$ cells | Range of mCherry Intensity amongst $CC3^{+}$ cells | Geometric Mean of mCherry Intensity amongst $CC3^{+}$ cells |
| Expt 1 | 1.5% | $1x10^{5}$-$1x10^{6}$ | $3x10^{5}$ |
| Expt 2 | 2.2% | $1x10^{5}$-$5x10^{6}$ | $4x10^{5}$ |
| Expt 3 | 5.1% | $1x10^{5}$-$3x10^{7}$ | $4x10^{5}$ |

| | WT 37°C-Infected Monolayers | |
|---|---|---|
| | Avg % of $CC3^{+}$ cells | # M cells analyzed per expt ([a]) |
| Expt 1 | 0.2% | 10 (1) |
| Expt 2 | 0.8% | 10 (0) |
| Expt 3 | 5.1% | 20 (3) |

[a] Number CC3+ M cells per experiment

Supplemental Table 4. Table of significances for Figure 6J.[a]

| | 0 *Yptb* Internalized | | | | |
|---|---|---|---|---|---|
| | WT 37°C | Δ*yopE* | *yopER144A* | *yopESptP* | *yopEL109A* |
| WT 37°C | | | | | |
| Δ*yopE* | **** | | | | |
| *yopER144A* | **** | ns | | | |
| *yopESptP* | ns | ** | ** | | |
| *yopEL109A* | ns | **** | **** | ns | |

| | 1-9 *Yptb* Internalized | | | | |
|---|---|---|---|---|---|
| | WT 37°C | Δ*yopE* | *yopER144A* | *yopESptP* | *yopEL109A* |
| WT 37°C | | | | | |
| Δ*yopE* | ns | | | | |
| *yopER144A* | ns | ns | | | |
| *yopESptP* | ns | ns | ns | | |
| *yopEL109A* | ns | ns | ns | ns | |

| | 10+ *Yptb* Internalized | | | | |
|---|---|---|---|---|---|
| | WT 37°C | Δ*yopE* | *yopER144A* | *yopESptP* | *yopEL109A* |
| WT 37°C | | | | | |
| Δ*yopE* | *** | | | | |
| *yopER144A* | **** | ns | | | |
| *yopESptP* | ns | ns | ns | | |
| *yopEL109A* | ns | ** | *** | ns | |

[a] ** $p < 0.01$; *** $p < 0.001$; **** $p < 0.0001$; ns = no significance

Supplemental Table 5. *Yersinia* strains used in this study.

| Strain Number | Description | Original Strain # | Reference for original strain |
|---|---|---|---|
| ACF053 | *Yptb* YPIII pIB1, pACYC184-ptet::GFP | JM301 | [1] |
| ACF054 | *Yptb* YPIII pIB1, Δ*yadA* + pACYC184-ptet::GFP | MM92 | this study |
| ACF055 | *Yptb* YPIII pIB1, Δ*inv* + pACYC184-ptet::GFP | ERG9 | this study |
| ACF056 | *Yptb* YPIII pIB1, Δ*inv/yadA* + pACYC184-ptet::GFP | FM420 | this study |
| ACF068 | *Yptb* YPIII pIB1, Δ*yopE/sycE*+pACYC184-ptet::GFP | LL31 | [1] |
| ACF073 | *Yptb* YPIII pIB1, Δ*yscN-U* +pACYC184-ptet::GFP | JMB31 | [2] |
| ACF076 | *Yptb* YPIII pIB1, pMMB67EH-ptetA::mCherry | JM301 | [1] |
| ACF085 | *Yptb* YPIII pIB1, Δ*yopE* + YopEL109A +pACYC184-GFP | WS315a | [3] |
| ACF087 | *Yptb* YPIII pIB1, Δ*yopE* + YopER144A + pACYC184-GFP | WS312 | [3] |
| ACF091 | *Yptb* YPIII pIB1, Δ*yopE* + YopE1-100SptP166-293 + pACYC184-ptet::GFP | WS352 | [3] |
| ACF093 | *Yptb* YPIII pIB1, Δ*yopE* + YopE recombined +pACYC184-ptet::GFP | WS338 | [3] |
| ACF112 | *Yptb* YPIII pIB1, Δ*yopH* + pACYC184-ptet::GFP | this study | this study |
| ACF113 | *Yptb* YPIII pIB1, Δ*yopEH* + pACYC184-ptet::GFP | this study | this study |
| ACF114 | *Yptb* YPIII pIB1, Δ*yopO* + pACYC184-ptet::GFP | this study | this study |
| ACF115 | *Yptb* YPIII pIB1, Δ*yopEO* + pACYC184-ptet::GFP | this study | this study |
